# *RLSuite*: An integrative R-loop bioinformatics framework

**DOI:** 10.1101/2022.07.13.499820

**Authors:** H. E. Miller, D. Montemayor, S. Levy, K. Sharma, B. Frost, A. J. R. Bishop

## Abstract

R-loops are three-stranded nucleic acid structures containing RNA:DNA hybrids. While R-loop mapping via high-throughput sequencing can reveal novel insight into R-loop biology, the analysis and quality control of these data is a non-trivial task for which few bioinformatic tools exist. Herein we describe *RLSuite*, an integrative R-loop bioinformatics framework for pre-processing, quality control, and downstream analysis of R-loop mapping data. RLSuite enables users to compare their data to hundreds of public datasets and generate a user-friendly analysis report for sharing with non-bioinformatician colleagues. Taken together, RLSuite is a novel analysis framework that should greatly benefit the emerging R-loop bioinformatics community.

## INTRODUCTION

R-loops are three-stranded nucleic acid structures comprised of an RNA:DNA hybrid and displaced single strand DNA. R-loops form during transcription as nascent RNA re-anneals to the template DNA strand, often in regions with high G/C-skew, such as CpG islands (1). When dysregulated, R-loops can promote collisions between the transcription and replication machinery, leading to a stalled replication fork and genome instability (2). Most R-loop research has focused on this pathological context, particularly regarding its relevance for cancer (3–5) and aging (6,7). There is, however, mounting evidence that R-loops do not contribute to genome instability under basal conditions. From a recent meta-analysis, we found that R-loops are associated with approximately 36.9% of expressed genes (calculation provided in the code which accompanies this work; see *Availability*) and cover approximately 4.3% of the entire genome (8). This includes enhancers and topologically-associated domain (TAD) boundaries (8), suggesting that R-loops may regulate genomic architecture and transcription, a notion supported by the direct interaction between R-loops and cohesin (9). Additionally, recent evidence has implicated R-loops in DNA repair (10), reprogramming to pluripotency (11), and a host of other genomic and epigenomic processes (12). While these findings suggest the existence and importance of physiological R-loops, the locations and dynamics of these R-loops remains poorly understood.

R-loop mapping techniques enable the genome-wide analysis of R-loop locations and dynamics. In 2012, *Ginno et al*. introduced the first R-loop mapping technique, DNA:RNA immunoprecipitation sequencing (DRIP-Seq) (13). In the decade since, more than 40 R-loop mapping studies have been described (8). While these studies have greatly expanded our understanding of R-loop dynamics, they have also revealed pervasive quality control issues that limit the interpretation of their findings (14). In recent work, we mined 810 public R-loop mapping datasets and developed a robust quality control (QC) approach for R-loop mapping data. With this QC approach, we identified high-quality R-loop datasets and performed a large-scale meta-analysis that revealed novel biological insights regarding the differences between *in situ* and *ex vivo* mapped R-loops (8). Moreover, we built a database of reprocessed and standardized R-loop mapping datasets, and described novel methods for analysing R-loop data (8,15). With the data and code developed in these studies, we created *RLSuite*, a collection of software packages for the analysis of R-loop mapping data.

There are a growing number of publications that describe bioinformatic approaches for the study of R-loops. However, most have focused on the task of predicting R-loop formation from genomic sequences (16–21) and associated epigenomic marks (22). At present, only two bioinformatic tools for the analysis of R-loop mapping data have been described. The first, *footLoop*, is designed specifically for the analysis of long-read bisulfite-converted R-loop mapping data (SMRF-Seq) (23), a type of sequencing that is not applicable for most R-loop mapping studies. The second, the DRIP-optimized Peak Annotator (DROPA), facilitates the annotation of DRIP-seq peaks using genomic features and gene expression information (24), but does not provide users with the capability to perform upstream data processing, quality control their data, or analyze and explore their data in the context of public R-loop mapping datasets.

To address this critical resource gap, we have developed RLPipes, RLHub, and RLSeq (collectively termed “RLSuite”). RLSuite provides an integrative workflow for R-loop data analysis, including automated pre-processing of R-loop mapping data using a standard pipeline, multiple robust methods for quality control, and a range of tools for the initial exploration of R-loop data. RLSuite also provides the unique capability to compare analysis results with those from the public R-loop datasets that we previously reprocessed (8,15). Moreover, it provides a user-friendly HTML analysis report which should prove useful for both bioinformaticians and biologists alike.

## RESULTS

In recent work, we reprocessed 810 R-loop mapping datasets, establishing a robust method of quality control (8). We then used these data to build *RLBase*, the largest R-loop mapping database yet described (15). We also established novel approaches for the analysis of R-loop mapping data, and demonstrated the utility of analysing R-loop mapping data in the context of publicly available R-loop datasets (8). In the present study, we codified and standardized our analysis approaches in the form of three software packages: *RLPipes, RLHub*, and *RLSeq*. RLPipes is a Bioconda command line tool for the automated processing of R-loop mapping data from raw files, BAM files, or public data accessions. RLHub is an R/Bioconductor “ExperimentHub” package that facilitates access to the processed datasets stored within RLBase. Finally, RLSeq is an R/Bioconductor package that provides a comprehensive toolkit for quality control, exploration, and visualization of R-loop mapping data. Collectively termed “RLSuite”, these packages provide the first integrative R-loop bioinformatics framework. In the following sections, we describe the workflow for analysis of R-loop mapping data with *RLSuite*.

### RLPipes: Upstream pre-processing of R-loop mapping data

A typical workflow for RLSuite involves providing raw data files (and relevant metadata) or public data accessions to RLPipes, which then automatically processes the data (**Figure 1A**). RLPipes is a command-line tool based on *Snakemake* (25) which implements a uniform workflow for data wrangling, QC, alignment, peak calling, and coverage analysis. This workflow follows standard practices for the analysis of genomic and epigenomic data (26). Finally, the resulting data is analysed with RLSeq to generate an analysis report (see *RLSeq: downstream R-loop data QC, analysis, and visualization*).

**Figure 1:**
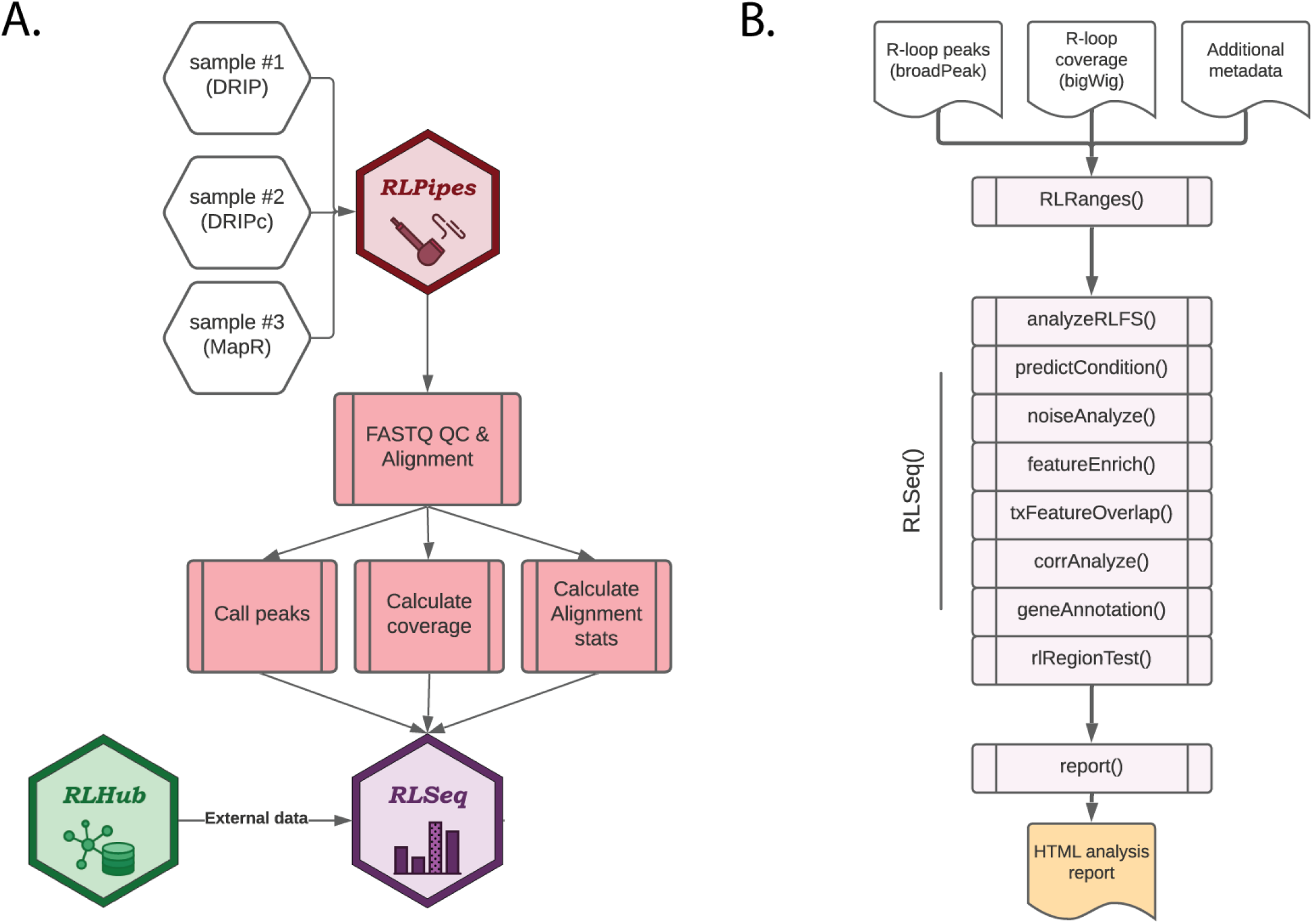
Overview of RLSuite workflow. (A) Flow diagram showing the upstream analysis of R-loop mapping data with *RLPipes*, terminating in *RLSeq* analysis (with supporting data from *RLHub*). Samples of nearly any R-loop mapping modality can be supplied to RLPipes either separately or in combination. The RLPipes workflow performs quality control on the raw read data and aligns it to the genome. Then, alignment statistics, coverage, and peaks are calculated. Finally, the RLSeq analysis workflow is implemented. (B) Flow diagram depicting the RLSeq analysis workflow. Input data are converted to an *RLRanges* object. In a simplified RLSeq analysis workflow, the *RLSeq()* function is invoked. This triggers all eight analysis steps, and the results are saved in the RLRanges object. Finally, the *report()* function is invoked to generate a user-friendly HTML analysis report.

To initiate an RLPipes workflow, a user can invoke the *RLPipes build* command. If supplying local files, RLPipes will use a subset of the data to infer parameters such as read length. If public data accessions are provided, then RLPipes will retrieve the sequence metadata from the Sequence Read Archive (SRA). After validating the input parameters, RLPipes generates a directory to house the workflow outputs along with a configuration file. The user is then prompted to invoke the *RLPipes check* command, launching a dry run of the workflow and generating an image file showing the proposed analysis workflow steps. Finally, the user invokes the *RLPipes run* command, which launches all workflow jobs, leading to the generation of BAM files, coverage tracks, peaks, and RLSeq outputs (described in “RLSeq”).

### RLHub: A hub for R-loop data

RLHub is an R/Bioconductor ExperimentHub (27) package that provides a convenient R interface for users to access processed datasets relevant for the analysis of R-loop mapping data. RLHub contains 16 datasets (**Table S1**), most of which are required for the functions in RLSeq. These datasets can be loaded via built-in accessor functions. For example, *rlbase_samples()* loads the manifest of all publicly-available R-loop mapping samples in *RLBase*.

Additionally, *rlbps()* loads the R-loop binding proteins identified from three recent studies (28–30), with their associated confidence scores. It also includes the machine learning models used for predicting R-loop mapping data quality (accessed via *fft_model()*) (8). When a dataset is accessed for the first time, it is downloaded to a cache on the user’s local machine to enable rapid access upon future invocations. In summary, RLHub provides a convenient R interface for accessing a wealth of processed data relevant for the analysis of R-loops.

### RLSeq: downstream R-loop data QC, analysis, and visualization

*RLSeq* is an R/Bioconductor package for the downstream analysis of R-loop mapping data. It is run automatically as part of the *RLPipes* workflow, but it can also be used independently in any standard R session. In a minimal RLSeq analysis workflow, a user will load their data as an *RLRanges* object with *RLRanges()*, run the *RLSeq()* function to perform core analysis steps with default parameters, and then run the *report()* function to generate a HTML analysis report which can be shared with colleagues (**Figure 1B**). In the following sections, we will describe each of the steps in this workflow. To illustrate the utility of *RLSeq*, we analysed DRIP-Sequencing datasets deposited by *Jangi et al*., *2017* as part of their study of the survival of motor neuron (SMN) gene (31). We analysed their DRIP-seq data from two SH-SY5Y cell lines, one with short hairpin RNA (shRNA) which is untargeted (shCTR; SRA ID: SRX2187024), and one with shRNA targeting SMN (shSMN; SRA ID: SRX2187025). The following sections will describe the utilities in *RLSeq* along with the results from running those functions on the SH-SY5Y shCTR dataset and, where indicated, the shSMN dataset as well.

#### RLRanges

The functions in *RLSeq* operate on a custom object class, “RLRanges.” This class is an extension of the *GenomicRanges* class from Bioconductor (32) and contains additional information specific to the analysis of R-loop data within RLSeq. RLRanges can be created using the *RLRanges()* function when providing coverage, peaks, and metadata (**Figure 1B**). Additionally, public datasets in RLBase can be loaded as RLRanges objects using the *RLRangesFromRLBase()* function. This facilitates access and analysis of public datasets within the RLSeq framework. For the present analysis, *RLRanges* for SH-SY5Y shCTR and shSMN were created using the *RLRanges()* function as part of the *RLBase* data generation workflow (see *Availability*).

#### RLFS analysis

The first step in the *RLSeq* workflow is to perform R-loop forming sequences (RLFS) analysis using the *analyzeRLFS()* function. RLFS are computationally-predicted genomic regions that show favourability for R-loop formation (17–20). In previous work, we demonstrated that high quality R-loop mapping datasets show robust enrichment in these regions, indicating their utility for predicting whether R-loop mapping was successful (8). In that work, we developed RLFS analysis, a method which employs permutation testing to evaluate the statistical enrichment of R-loop mapping peaks within RLFS (8). Within RLSeq, this analysis is performed via the *analyzeRLFS()* function, and the results of the analysis are visualized with the *plotRLFSRes()* function (**Figure 2**). In the present study, we ran RLFS analysis on SH-SY5Y shCTR DRIP-seq data and visualized the results (**Figure 2**). From the resulting plot, we observed that the sample peaks were strongly enriched within RLFS, suggesting the data represents successful R-loop mapping.

**Figure 2:**
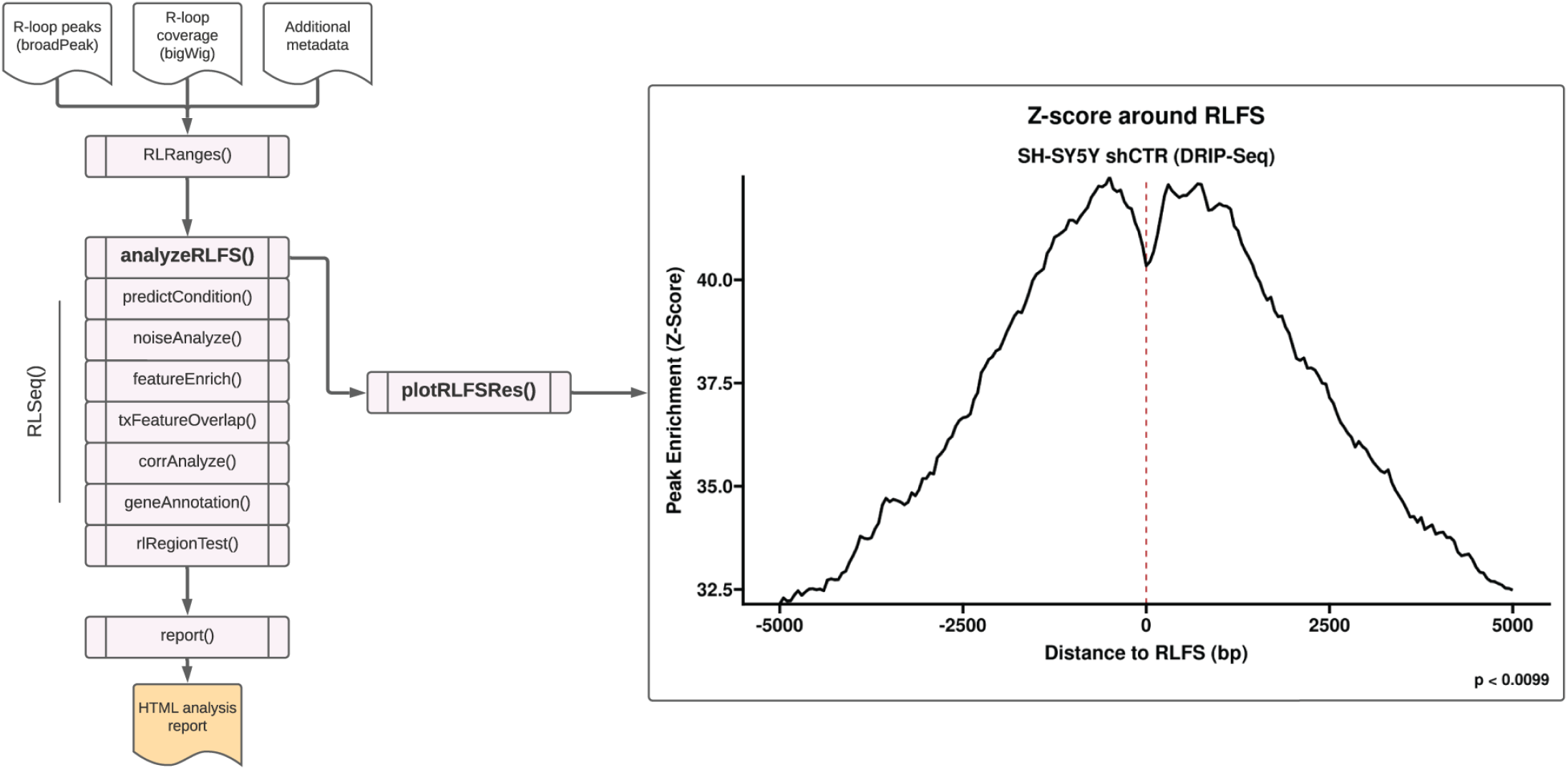
R-loop forming sequences (RLFS) analysis. RLFS analysis was applied to analyze publicly available DRIP-seq data from SH-SY5Y cells with control shRNA (“shCTR”) (SRA ID: SRX2187024). This analysis was invoked with the *analyzeRLFS()* function and plotted with the *plotRLFSRes()* function. The analysis involved permutation testing to determine the statistical enrichment of user-supplied R-loop peaks within computationally predicted RLFS. The RLFS analysis plot displays the p value from permutation testing (with the standard 100 permutations, the minimum possible p value is 0.0099). Additionally, the peak enrichment around RLFS is displayed as a Z score.

#### Sample quality classification

As demonstrated in our previous work, R-loop mapping data is prone to quality issues which can severely limit interpretability (8). In that work, we developed a machine learning model which uses the results of RLFS analysis to predict whether a dataset shows evidence of successful R-loop mapping (8). Samples that are predicted to map R-loops successfully receive a model prediction of “POS” (positive), whereas samples that are predicted to not map R-loops get a prediction of “NEG” (negative). The prediction is performed using the *predictCondition()* function. The prediction function also generates a list object which includes the normalized model predictors for the sample, along with the four criteria required for receiving a “POS” prediction (see *Methods*). For the sample analysed in this example (SH-SY5Y shCTR), the result of this prediction was “POS,” indicating that it robustly maps R-loops as expected.

#### Noise analysis

As with all QC methods, the method implemented by *predictCondition()* has limitations (8) and a comprehensive QC workflow should include additional complementary methods. Towards that end, RLSeq includes multiple additional QC methods. One such approach, termed “noise analysis” (invoked via the *noiseAnalyze()* function), quantifies the signal-to-noise ratio of R-loop mapping coverage using the method developed by *Diaz et al* (33). Briefly, random 1kb genomic bins are created and the R-loop mapping coverage signal is quantified within each bin. For high-quality datasets, it is expected that most signal will be found within a relatively small subset of bins, as this indicates strong enrichment of signal within specific genomic regions (as would be expected for successful R-loop mapping). The results are visualized qualitatively using a “fingerprint plot” (**Figure 3A**) (named after the deepTools implementation of this same approach (34)). Created with the *plotFingerprint()* function, these plots reveal the proportion of signal captured by a certain proportion of genomic bins. A “hockey-stick” shape (greater inflection) is a sign of good signal to noise ratio because it implies that most signal is found within a subset of the genome (as would be expected for R-loop mapping). From our analysis of the SH-SY5Y shCTR DRIP-seq sample, we observed a strong “hockey-stick” pattern (**Figure 3A**). This further suggests the high quality of that dataset.

**Figure 3:**
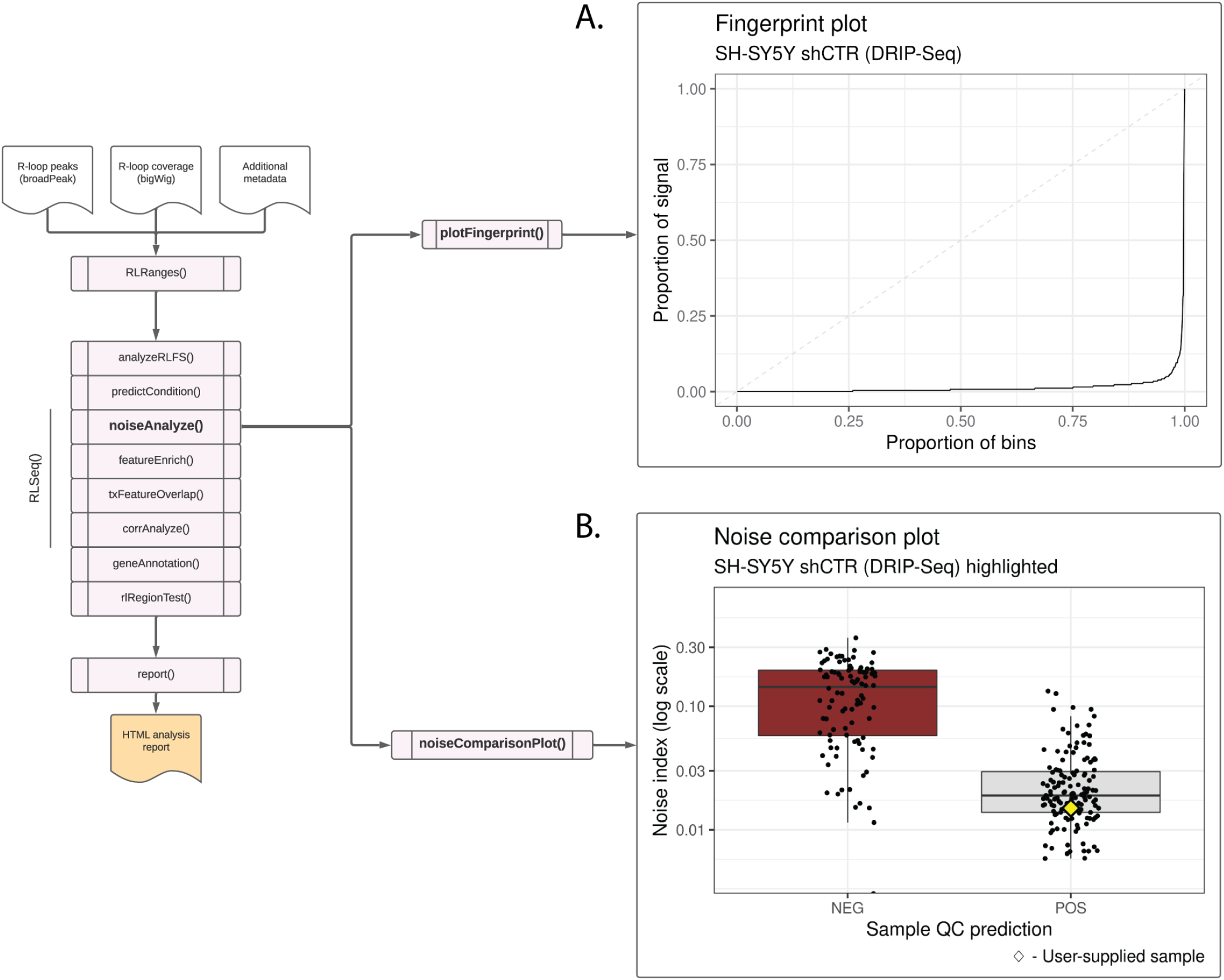
Noise analysis of SH-SY5Y DRIP-seq data (SRX2187024). (A) A fingerprint plot showing the proportion of signal contained in genomic bins. The strong inflection implies that most signal is contained in relatively few bins, indicating a good signal-to-noise ratio. This plot was generated using the *plotFingerprint()* function. (B) A noise comparison plot showing the noise index for DRIP-Seq samples in the RLBase database. The plot shows that the noise index of the user-supplied sample (SH-S5Y5 shCTR; represented by a yellow diamond) is low, further suggesting good signal-to-noise ratio. This plot was generated via the *noiseComparisonPlot()* function.

*RLSeq* provides the capability to compare noise analysis results from a user-supplied sample with those from publicly available datasets of the same R-loop mapping modality. This plot, termed a “noise comparison plot” and generated via the *noiseComparisonPlot()* function, displays the average standardized noise (“Noise index”) for public DRIP-seq samples alongside the user-supplied sample (**Figure 3B**). Samples are separated based on their “prediction” (from the quality model). This plot enables users to not only quantify the noise within a sample, but also see how it compares to similar samples from previous studies. From the analysis of the SH-SY5Y sample, we observe that it has a relatively low noise index, like other “POS”-predicted DRIP-seq samples (**Figure 3B**). This further indicates the high quality of this dataset.

#### Feature enrichment test

The “feature enrichment test” (invoked via the *featureEnrich()* function), analyses the enrichment of user-supplied peaks within various genomic features. This test can be used as a QC method because high-quality R-loop mapping peaks are statistically enriched within certain genomic features, including gene body features (e.g., exons), CpG islands, G4 quadruplexes, and regions of high G/C-skew (8). Beyond providing a test of data quality, this analysis also enables the exploration of peak enrichment within a host of genomic features, including repetitive elements, transcription factor binding sites, and cis-regulatory elements. Moreover, by comparing multiple samples analysed with *featureEnrich()*, users can gain initial insight into the differences between the types of R-loops mapped across experimental conditions. Once analysis is complete, the data is visualized using the *plotEnrichment()* function (**Figure 4A**). Like previous approaches, this visualization enables users to view their results in the context of similar samples from public datasets and determine whether their results diverge from previous datasets.

**Figure 4:**
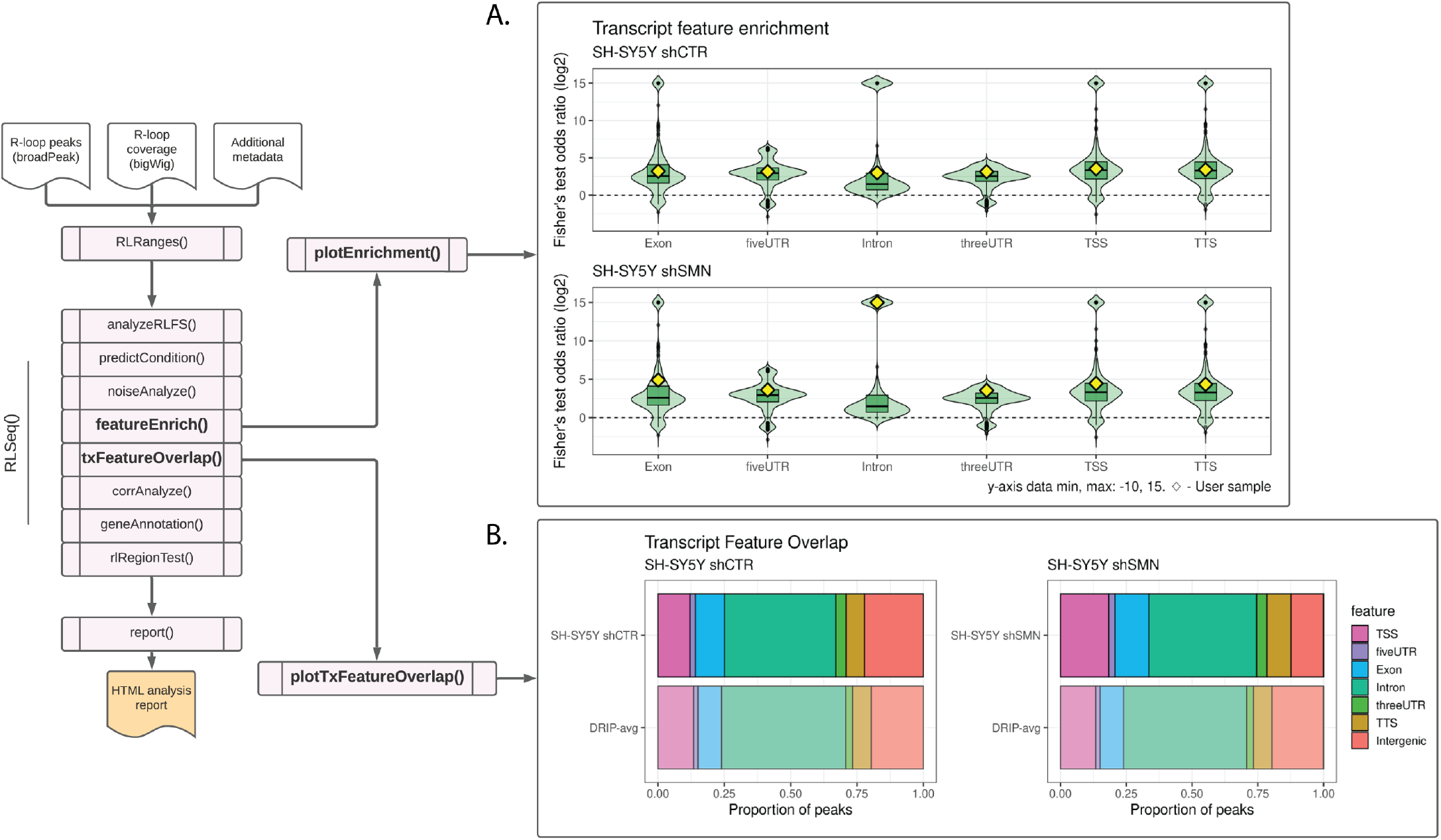
Feature analysis results in SH-SY5Y DRIP-seq data from shCTR (SRX2187024) and shSMN (SRX2187025) cells. (A) A feature plot showing the enrichment of RLBase samples within various transcript features. The Y-axis is the odds ratio (log2 transformed) from Fisher’s exact test. The diamonds represent the odds ratios for the user-supplied sample. The plot was generated using the *plotEnrichment()* function. (B) A transcript feature overlap plot that shows the proportion of user-supplied peaks overlapping transcript features. The average of all high-quality DRIP-Seq datasets is also shown for comparison. The plot was generated using the *plotTxFeatureOverlap()* function.

For the example analysis in this work, we selected two DRIP-seq datasets upon which to perform feature enrichment, SH-SY5Y shCTR and SH-SY5Y shSMN (SMN knock-down) cells. The study from which these data are derived reports that knock-down of SMN (shSMN) causes an increase in intron retention (31). Moreover, they observed an increase in R-loop formation within gene body features, specifically retained introns and exons. From our analysis of their shCTR and shSMN samples, we likewise observed an overall increase in enrichment within transcriptomic features, and particularly within introns (**Figure 4A**), thus recapitulating the findings from that study.

#### Transcript feature overlap

In addition to the more rigorous statistical analysis described above, it is also useful to broadly analyze the proportion of R-loop mapping peaks which overlap various transcriptomic features, such as exons and the transcription start site (TSS). To facilitate this type of analysis, *RLSeq* provides the *txFeatureOverlap()* function. This function finds the overlap of user-supplied peaks with transcript features and summarizes them to determine the proportion of peaks overlapping each type of feature. Of note, similar to other types of analysis, *txFeatureOverlap()* uses a “feature priority” to uniquely assign peaks to a single feature, a step that is required when peaks overlap multiple features (see details in *Methods*). Finally, the results are then visualized with the *plotTxFeatureOverlap()* function. This visualization displays the proportion of peaks from the user-supplied sample overlapping various features (**Figure 4B**).

Like the Feature enrichment test analysis above, we applied the Transcript feature overlap analysis to both SH-SY5Y shCTR and shSMN samples. Similar to the prior analysis, we successfully recapitulated the increase in overall gene body R-loops (**Figure 4B**). While we did not observe a strong increase in introns, this is not unexpected given that this style of analysis enforces a feature priority which disadvantages introns (see *Methods*). However, we observed an increase in Exon and TSS (**Figure 4B**), recapitulating the prior study’s findings (31). In general, RLSeq is not designed for differential analysis of R-loop locations between conditions. However, the findings from the above analyses show that RLSeq is useful for initial exploration of quality and genomic feature enrichment across samples and conditions. They even recapitulated findings from the previous publication regarding the impact of SMN knock-down on R-loop localization (31). However, deeper insights can be gained from more robust statistical approaches, and, in future work, we plan to introduce tools specifically designed for the differential analysis of R-loop abundance across conditions.

#### Correlation analysis

Correlation analysis (invoked via the *corrAnalyze()* function) is an additional quality control methodology, originally described in the recent work of *Chedin et al* (14). The approach involves calculating the pairwise correlation of coverage signal between public R-loop mapping samples and a user-supplied sample (14). The signal correlation is calculated in the genomic regions surrounding “gold-standard R-loops” (R-loops that were uncovered by single molecule, long-read, bisulfite-based R-loop mapping (SMRF-Seq) (23)). The results are visualized in a correlation heatmap, created with *corrHeatmap()*. This visualization provides a detailed picture of how well samples agree with one-another and how well a user-supplied sample agrees with similar previous datasets. We applied the correlation analysis method to the SH-SY5Y shCTR DRIP-seq data and plotted the resulting heatmap (**Figure 5**). We observed that the shCTR SH-SY5Y dataset clustered with similar positive DRIP-seq samples (**Figure 5**), further suggesting that the dataset is of high quality.

**Figure 5:**
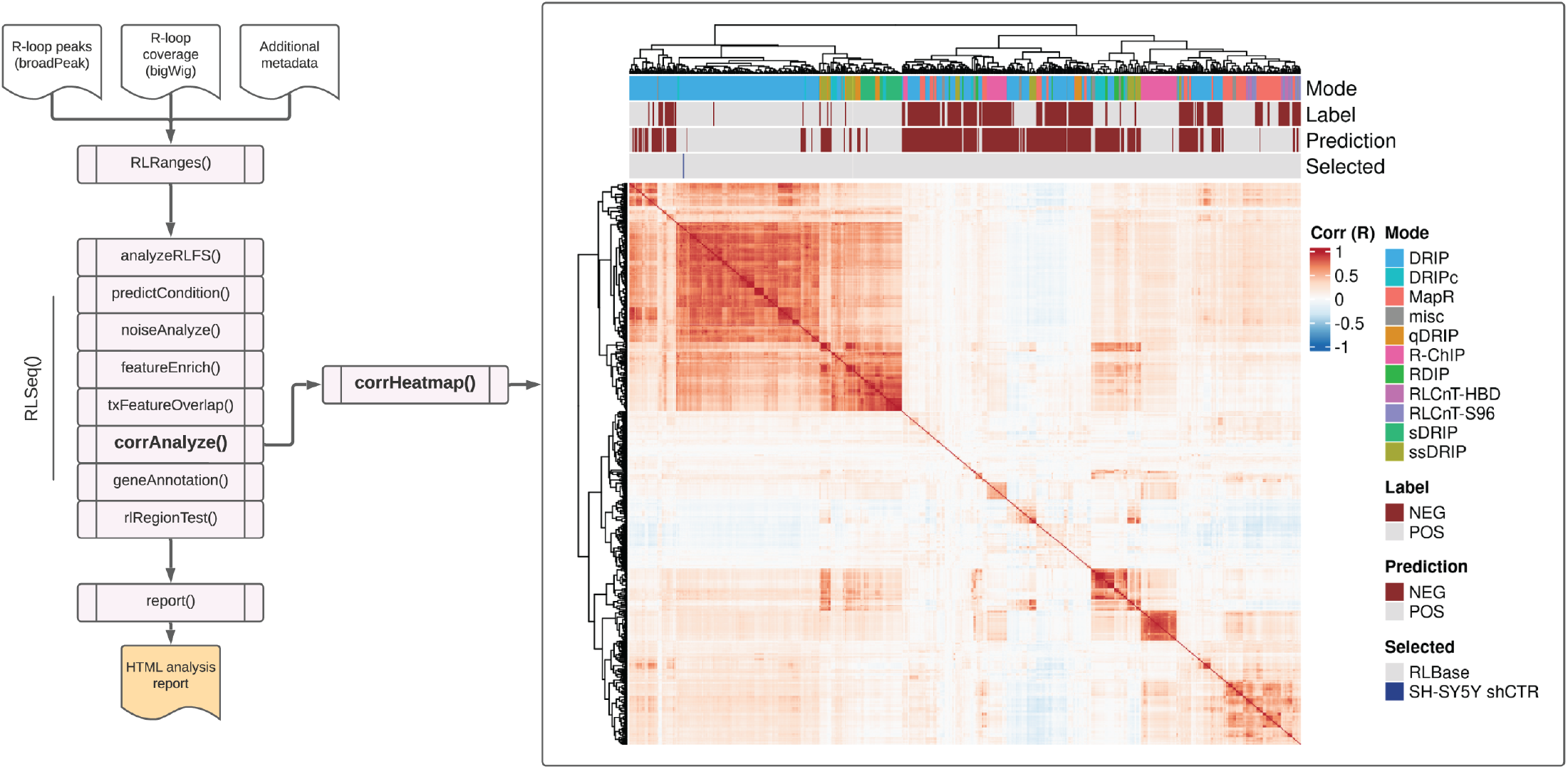
Correlation analysis results from SH-SY5Y DRIP-seq data (SRX2187024). A correlation heatmap showing the pairwise Pearson correlation between the human samples in the RLBase database around “gold-standard R-loops” (see *Methods*). The location of the user-supplied sample is shown in the “Selected” annotation row on the top of the plot. The plot was generated using the *corrHeatmap()* function.

#### Gene annotation

The gene annotation feature provides a convenient function for identifying the genes with which the user-supplied R-loop mapping peaks overlap. The analysis is performed via the *geneAnnotation()* function and yields a table of peaks and corresponding gene IDs. We applied this analysis to the SH-SY5Y shCTR data and returned the overlapping genes for each peak (**Table S1**).

#### RL-Regions test

Finally, *RLSeq* provides the capability to analyze the overlap between user-supplied peaks and R-loop regions (RL-Regions), regions of consensus R-loop formation uncovered in our previous work (8), via the *rlRegionTest()* function. These regions represent areas of the genome which show conserved R-loop formation across studies and samples. The degree to which the user-supplied peaks overlap these regions is a useful measure of quality. It also indicates the degree to which the user-supplied peaks detect non-canonical R-loops (those not typically detected in mapping experiments). The data are visualized as a Venn diagram using the *plotRLRegionOverlap()* function (**Figure 6**). From analysis of the shCTR SH-SY5Y DRIP-seq data, we observe a strong overlap with “canonical” regions of R-loop formation (**Figure 6**).

**Figure 6:**
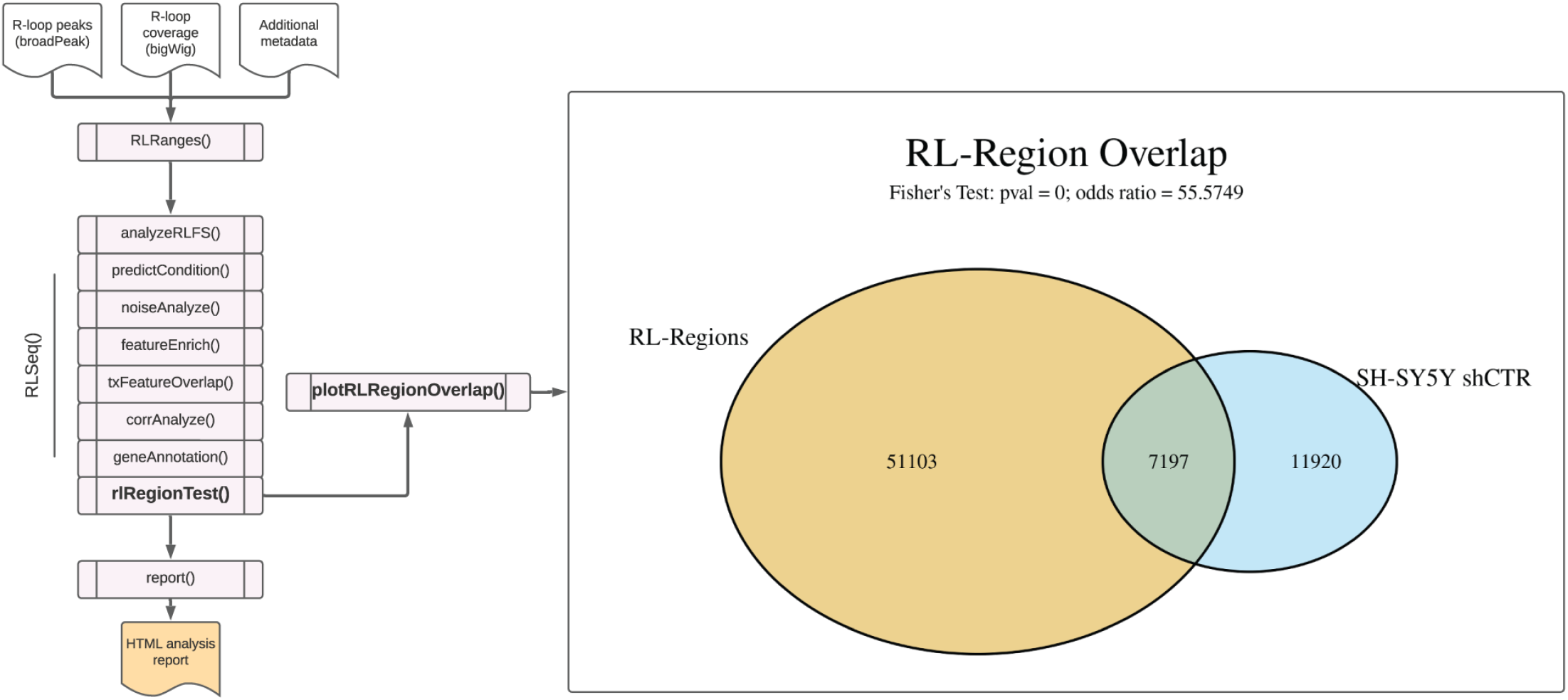
RL Region analysis results from SH-SY5Y DRIP-seq data (SRX2187024). A Venn diagram showing the overlap of user-supplied R-loop peaks with R-loop Regions (RL Regions), regions of conserved R-loop formation (8). This plot was generated via the *plotRLRegionOverlap()* function.

#### The RLSeq workflow

The analyses described above represent the core analyses available within the *RLSeq* R/Bioconductor package. While many bioinformaticians will be sufficiently skilled in R programming to easily navigate and use the *RLSeq* package as part of their preferred workflow, biologists and novices in R programming may find greater difficulty. Therefore, we have provided a convenient wrapper, *RLSeq()*, which automatically invokes the above functions (**Figure 1B**).

#### The RLSeq analysis report

Finally, we provide the *report()* function, which generates a user-friendly HTML analysis report for sharing results with colleagues (**Figure 7**). The report begins with a summary of the analysis run, including relevant metadata describing the sample and software version, and a table summarizing the available results. The report then shows all the analysis steps and visualizations along with detailed descriptions of the methods used to generate those results. Additionally, interactive data tables are provided to facilitate the exploration of results, and download buttons are enabled which provide the capability to download tabular data from the report in comma-separated values (CSV) format. Overall, this report provides a valuable interface for non-bioinformaticians to understand the results from *RLSeq*.

**Figure 7:**
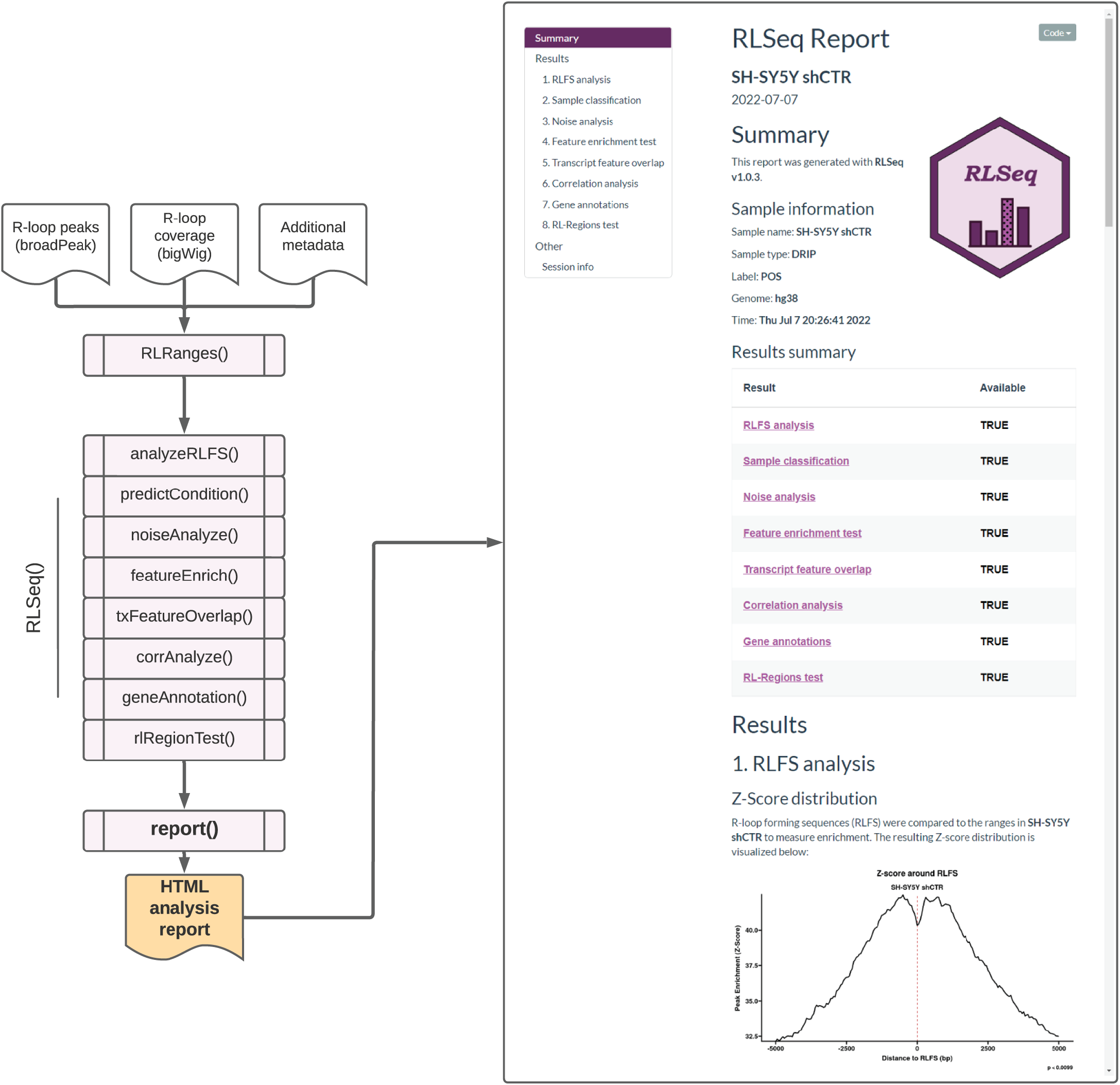
Screenshots of the RLSeq analysis report for SH-SY5Y DRIP-seq data (SRX2187024). The report HTML file was generated by running *report()* and then opened in a web browser. The figure depicts a screenshot of the overview section of the RLSeq report. This section shows the metadata from the analysis along with the list results available within the report. It also shows the RLFS analysis page, which reveals the results from permutation testing. A link to the full report is provided within this work (see *Availability*).

To illustrate the utility of the HTML analysis report, we executed the *report()* function with the SH-SY5Y shCTR dataset. The resulting report displays the analysis results (shown in previous sections) and includes detailed methods descriptions along with interactive tables that can be downloaded in CSV format (**Figure 7**). A web URL for this report is also provided herein (see *Availability*).

## DISCUSSION

In the present study we describe *RLSuite*, a collection of three software packages (*RLPipes, RLHub*, and *RLSeq*) that provides an integrative workflow for R-loop data analysis. RLSuite provides users with the capability to easily process, quality control, and explore their data. Moreover, the workflow produces user-friendly HTML reports which facilitate sharing results with non-bioinformatician colleagues. Finally, by leveraging the wealth of processed data developed previously (accessible via the RLHub package), RLSeq provides users with the capability to both explore their own data and compare their results to those obtained from publicly available samples, thus providing useful context for interpreting the insights gained from the analysis. It also offers a limited capability to compare results between samples from different biological conditions, as we demonstrated in our analysis of SH-SY5Y DRIP-seq data (**Figure 4**). However, a full and robust differential R-loop analysis will require additional tools not currently provided by RLSeq, and we intend for those features to be released in future versions of RLSuite.

At present, few studies have addressed the lack of R-loop-specific bioinformatics software packages. In 2019, *Russo et al*. introduced “DRIP-seq optimized peak annotator” (DROPA), a useful package for annotating DRIP-seq peaks and inferring strand assignment (24). However, DROPA does not provide quality control tools, the ability to compare a user-supplied sample to public datasets, nor the ability to examine peak enrichment across a wide array of annotations, as RLSeq does. It also does not provide the ability to perform upstream analysis of R-loop mapping datasets, as RLPipes does. Another package, *footLoop*, was developed by *Malig et al*. for upstream processing of long-read bisulfite-converted R-loop mapping data (SMRF-Seq) (23). However, *footLoop* is not applicable for typical R-loop mapping approaches, as these generally do not involve long-read sequencing or bisulfite foot printing. While there are additional tools which are useful for predicting genome sequences that are favourable for R-loop formation, particularly Qm-RLFS-finder (17) and SkewR (13), they do not enable the upstream or downstream analysis of R-loop datasets. RLSuite thus represents the first integrative R-loop bioinformatics framework, one which should prove valuable to the field. As new R-loop datasets become available and new features are implemented, we anticipate that the utility of RLSuite will only increase throughout the coming years.

## CONCLUSIONS

Herein we describe *RLSuite*, an integrative bioinformatics framework for the analysis of R-loop mapping data. RLSuite comprises three primary tools: *RLPipes* (an automated workflow for upstream processing of R-loop mapping datasets), *RLHub* (a hub for accessing processed R-loop datasets), and *RLSeq* (a comprehensive toolkit for the downstream analysis of R-loop mapping data). Taken together, RLSuite (RLPipes, RLSeq, and RLHub) provides a robust workflow for processing, quality control, and downstream analysis of R-loop mapping data. All three packages are accessible from open-source software repositories (Bioconda and Bioconductor) and the source code is available on GitHub (see *Availability*).

## METHODS

### Preliminary data curation

Prior to data curation, we gathered basic genome information and genomic features. This included: (1) UCSC genome information, (2) R-loop forming sequences (RLFS) predictions, and (3) genomic annotations tailored for R-loop analysis. The following subsections detail the processing steps involved in obtaining these data.

#### Genome metadata

Prior to running the data generation pipeline, it was necessary to curate a list of all available genomes in the UCSC data repository. For each genome in UCSC, the following information was obtained via the *makeAvailableGenomes*.*R* script in the *RLBase-data* GitHub repository (See *Code Availability*):

1. UCSC organism assembly ID
2. Taxonomy ID
3. Scientific Name
4. Year that the genome assembly was introduced
5. Genome length
6. Gene annotation availability (TRUE/FALSE)
7. Effective genome size at various read lengths (calculated via the *Khmer* python package). The resulting data was then packaged for use with *RLPipes* and *RLSeq*.

#### R-loop forming sequences (RLFS)

R-loop forming sequences were generated via methods we described previously (8).

#### Genomic annotation database

We developed a custom genomic annotation database for use with *RLSeq* that contained annotations relevant to R-loop biology. This was accomplished via a custom R script, *getGenomicFeatures*.*R*, in the *RLBase-data* GitHub repository (see *Code Availability*).

### RLPipes

RLPipes is a command-line interface (CLI) tool written in python and hosted on Bioconda (35) which is used for upstream processing of R-loop mapping data (see *Code availability*). It can accept FASTQ and BAM files, and it can also be used with Sequence Read Archive (SRA) or Gene Expression Omnibus (GEO) accessions. The typical workflow for using *RLPipes* follows the *build, check, run* pattern.

#### Build

*RLPipes build* is a command which processes a “sample sheet” into a configuration file which can be used directly with the underlying workflow. When used with local files (FASTQ or BAM), the *build* command simply verifies that the files all exist, that all the supplied arguments and options are valid, and it finds the library type (single-end or paired-end) and read length for each sample (using the *pysam* package (36) for BAM files and the *pyfastx* package for FASTQ files (37)).

If the samples supplied are SRA or GEO accessions, then the *pysradb* python package (38) is implemented to query the SRA database and return the following:

1. The SRA experiment accession
2. The SRA study accession
3. The experiment title (author-submitted)
4. The organism taxonomy ID
5. The SRA run accession(s)
6. The library layout (i.e., paired-end or single-end sequencing)
7. The total number of bases in the run
8. The total number of spots in the run

The organism taxonomy ID is then used to query the available genomes list (see *Preliminary data preparation*) and return the most recent genome assembly for the organism to be added to the standardized catalogue. Finally, the read length is calculated via floor division:

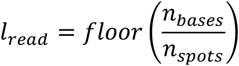

Where *L*_*read*_ is the read length, *n*_*bases*_ is the number of bases, and *n*_*spots*_is the number of spots in the sequencing run. Finally, the configuration is saved for use with downstream functions.

#### Check

The *check* operation in *RLPipes* is a useful tool for verifying the configuration with the underlying *snakemake w*orkflow manager (25) (equivalent of using the *dry-run* and *dag* flags in *snakemake*). It also provides a visualization depicting all the jobs which will be run as part of the workflow.

#### Run

All steps in the processing pipeline are run automatically via the *Snakemake* workflow manager (25) using pre-packaged *conda* environments. A typical analysis workflow will involve the following procedure: First, raw reads in SRA format are downloaded for each SRA run via the *prefetch* software from NCBI *sra-tools* (39). Then, reads are converted to “FASTQ” format using *fastq-dump* from *sra-tools* (39). Next, technical replicates are merged and interleaved (in the case of paired-end data) using *reformat*.*sh* from *bbtools* (40). Then, reads are trimmed and filtered with *fastp* (41), generating a quality report.

For R-loop mapping data, reads are aligned to the appropriate genome using *bwa mem* (42) or *bwa-mem2* (43), a faster implementation that users can enable with the *bwamem2* flag. Then, alignments are filtered (minimum quality score of 10 enforced), sorted, and indexed using *samtools* (44) and duplicates are marked using *samblaster* (45). Then, peaks are called using *macs3* (46) and coverage is calculated using *deepTools* (47). Optionally, *RLSeq* is then called by *RLPipes* to automatically perform downstream analysis steps (see description below).

### RLHub

RLHub is an *ExperimentHub* (27) R/Bioconductor package which provides convenient access to the processed datasets available in *RLBase*. It is available in Bioconductor v3.15 and has built-in accessor functions for quickly downloading and caching data. To generate these datasets, the *prepRLHub*.*R* script from the *RLBase-data* GitHub repository was executed (see *Code availability*). Full documentation of all objects and accessor functions provided through *RLHub* is provided by the *RLHub* reference manual (see *Code Availability*).

### RLSeq

*RLSeq* is an R/Bioconductor package for downstream analysis of R-loop mapping samples. It is available via Bioconductor v3.15 and it depends upon the *RLHub* package. *RLSeq* contains three primary functions: (1) *RLRanges*,(2) *RLSeq*, (3) *report*. The following sections detail the functions in the *RLSeq* package.

#### RLRanges

The primary class structure used in *RLSeq* is the *RLRanges* object, initialized with the *RLRanges* function. This object is an extension of the *GRanges* class from the *GenomicRanges* R package (32) to provide additional validation and storage for the following slots:

1. Peaks (the *GRanges* containing R-loop mapping peaks)
2. Coverage (a URL or file path to a bigWig file)
3. Mode (the type of R-loop mapping)
4. Label (“POS” or “NEG”, the author-assigned data label)
5. Genome (the UCSC genome ID)
6. Sample name (A sample name used for visualization and reporting)
7. *RLResults* (a list-like class for storing the results of *RLSeq* analysis)

#### Analyze R-loop forming sequences (*analyzeRLFS*)

R-loop forming sequences are regions of the genome with sequences that are favourable for R-loop formation (20). They are computationally predicted with the *QmRLFS-finder*.*py* software program (17) and serve as a test of whether a sample has mapped R-loops (8). The *analyzeRLFS* function provides a simple permutation testing method for analysing the enrichment of RLFS within a provided peakset. The full analysis procedure is described in detail in our recent work (8).

#### Predicting sample condition (*predictCondition*)

Following R-loop forming sequences (RLFS) analysis, the quality model is implemented for predicting the sample condition (i.e., “POS” if the sample robustly mapped R-loops and “NEG” if the sample resembles a negative control). This is accomplished with the *predictCondition* function which performs all the steps described in our recent work (8) to render a prediction for each sample. The results of this prediction, along with associated features and metadata, are stored in the *RLResults-predictRes* slot within the *RLRanges* object and returned to the user. For more detail, see the *RLSeq* reference manual (see *Code availability*).

#### Analyze noise profile (*noiseAnalyze*)

The *noiseAnalyze* function implements a modified version of the noise analysis method described by *Diaz et al* (33). Briefly, 1000 random genomic bins of 1Kb in size were generated used the *valr::bed_random()* function. For a query sample, the coverage is summed within all bins, standardized to the highest bin value. This step was performed for all public datasets in *RLBase*.

#### Feature enrichment testing (*featEnrich*)

A custom list of R-loop relevant genomic annotations was curated for the human (hg38) and mouse (mm10) genomes (see *Genomic annotations*) and made available via the *RLHub* R/Bioconductor package (see *RLHub annotations*). In *RLSeq*, each annotation type is tested for enrichment within supplied *RLRanges*, yielding enrichment statistics. The procedure for this testing is described in detail our recent work (8).

#### Transcript feature overlap analysis (*txFeatureOverlap*)

The *txFeatureOverlap* function performs a simplistic overlap analysis via the following procedure: (1) peaks are overlapped with transcriptomic features obtained from *RLHub* using the *bed_intersect* function from the *valr* R package (48). (2) For each peak which overlapped multiple features, a feature priority was used to assign it uniquely to a feature. The priority order is “TSS”, “TTS”, “fiveUTR”, “threeUTR”, “Exon”, “Intron”, and then “Intergenic”. For example, if a peak overlaps “TSS” and “Exon”, it will be uniquely assigned to “TSS”. (3) Once all peaks were uniquely assigned, they were saved as a table within the RLRanges object.

#### Correlation analysis (corrAnalyze)

The *corrAnalyze* function performs a sample-level correlation test that can be used to assess sample-sample similarity by calculating coverage signal (from genomic alignments) around high-confidence R-loop sites (14). The *corrAnalyze* function ingests an *RLRanges* object (with a valid *coverage* slot) and performs the following procedure: (1) the coverage is quantified within the high-confidence sites and added as a column to the signal matrix (see *gs_signal* reference in the *RLHub* documentation to learn more about this matrix), and (2) then the *cor* function in R is used to calculate the Pearson correlation to yield a correlation matrix. The correlation matrix is saved in the *RLResults-correlationMat* slot of the *RLRanges* object and returned to the user.

#### Gene annotation (geneAnnotation)

The *geneAnnotation* function provides a simple procedure for annotating *RLRanges* peaks with gene IDs by overlap. Briefly, gene annotations are automatically downloaded using the *AnnotationHub* R package (49) and then overlapped with the ranges in the *RLRanges* object using *bed_intersect* from *valr* (48). The mapping between peak IDs and gene IDs is saved in the *RLResults-geneAnnoRes* slot of the *RLRanges* object.

#### R-loop regions overlap test (*rlRegionTest*)

R-loop regions (RL regions) are R-loop consensus sites generated during the *RLBase-data* workflow (see *R-loop Regions*). The *rlRegionTest* function uses a simple procedure for finding RL regions which overlap with peaks in the supplied *RLRanges* object. It also calculates the significance and odds ratio of the overlap using Fisher’s exact test implemented via the *bed_fisher* function from the *valr* R package (48). The results of this test are saved in the *RLResults-rlRegionRes* slot of the *RLRanges* object.

#### RLSeq

The primary workflow in the *RLSeq* package can be conveniently run in one step with the *RLSeq* function. This command will run, in order, (1) *analyzeRLFS*, (2) *predictCondition*, (3) *noiseAnalyze*, (4) *featureEnrich*, (5) *txFeatureOverlap*, (6) *corrAnalyze*, (7) *geneAnnotation*, (8) *rlRegionTest*. The results are saved in the corresponding *RLResults* slots of the *RLRanges* object and returned to the user.

#### *RLSeq* plotting functions

*RLSeq* provides a variety of useful plotting functions which summarize and present analysis results:

*corrHeatmap*: generates a heatmap of the correlation matrix and annotations produced by *corrAnalyze* using either the *pheatmap* (50) or *ComplexHeatmap* (51) R packages.

*plotTxFeatureOverlap:* generates a stacked bar plot showing the proportion of user-supplied peaks assigned to various transcriptomic features by the *txFeatureOverlap* function. In addition, it also shows the average results of running *txFeatureOverlap* on high-quality, publicly available R-loop mapping samples generated from the same modality as the user-supplied sample.

*plotFingerprint:* generates a “fingerprint plot” (based on that which was developed by *Diaz et al* (33) and also implemented in deepTools (34)). The plot displays the results of running the *noiseAnalyze* function. Specifically, the standardized proportion of maximum bin signal was ranked, and the proportion of bins is plotted against it. Higher signal to noise ratio is indicated by the degree of inflection in the plot (i.e., most signal is contained within relatively fewer bins).

*noiseComparisonPlot:* generates a jitter+box plot showing the average max-standardized bin signal from *noiseAnalyze* in the user-supplied sample and the RLBase samples of the same modality. In this visualization, lower average signal across bins indicates better signal-to-noise ratio.

*plotEnrichment:* generates hybrid violin-box plots showing the Fisher’s exact test odds ratio for each annotation calculated in *featureEnrich*. Importantly, it displays these results for the user-supplied sample alongside the distribution of results for the public samples within the *RLBase* database.

*plotRLRegionOverlap:* generates a Venn diagram showing the overlap between the user-supplied sample peaks and the R-loop regions, along with overlap statistics as calculated by the *rlRegionTest* function.

*plotRLFSRes:* plots the Z-score distribution or Fourier transform of the Z-score distribution in a metaplot, based on the analysis results from *analyzeRLFS*.

#### Report

The *RLSeq report* function ingests an *RLRanges* object which has already been processed by the analysis functions in *RLSeq*. It then uses RMarkdown templates to automatically build a user-friendly HTML report showcasing all results with summary tables and plots (see the *RLSeq* reference for an example HTML report).

## ABBREVIATIONS

DRIP-Seq: DNA:RNA immunoprecipitation sequencing
QC: quality control
SRA: Sequence read archive
RLFS: R-loop forming sequences
TSS: Transcription start site

## DECLARATIONS

### Ethics approval and consent to participate

Not applicable

### Consent for publication

Not applicable

## Availability of data and materials

The scripts used to generate the figures in this study are available from GitHub: https://github.com/Bishop-Laboratory/RLSuite-Paper-Miller-2022

The HTML analysis report for shCTR SH-SY5Y is hosted on AWS at the following URL: https://rlbase-data.s3.amazonaws.com/misc/sh-sy5yrlseq-report.html

The datasets generated and/or analysed during the current study are available from the public RLBase AWS repository: s3://rlbase-data/ and https://rlbase-data.s3.amazonaws.com/. The scripts used to generate these data are found in the *RLBase-data* GitHub repository: https://github.com/Bishop-Laboratory/RLBase-data.

The data can be accessed via the AWS command line interface, the RLHub Bioconductor package (https://bioconductor.org/packages/release/data/experiment/html/RLHub.html), or the RLBase web database (https://gccri.bishop-lab.uthscsa.edu/rlbase/).

*RLPipes* is accessible at https://anaconda.org/bioconda/rlpipes - source code: https://github.com/Bishop-Laboratory/RLPipes. *RLHub* is accessible at https://bioconductor.org/packages/release/data/experiment/html/RLHub.html - source code: https://github.com/Bishop-Laboratory/RLHub. *RLSeq* is accessible at https://bioconductor.org/packages/release/bioc/html/RLSeq.html - source code: https://github.com/Bishop-Laboratory/RLSeq.

## Competing interests

None declared.

## Funding

NIH/NCI [R01CA152063 and 1R01CA241554], CPRIT [RP150445] and SU2C-CRUK [RT6187] to A.J.R.B and Greehey Graduate Fellowship Award and NIH/NIA [F31AG072902] to H.E.M. DOD [CDMRP PR181598] to K.S.

## Authors’ contributions

HEM and AJRB designed the study. HEM was the primary developer on all software and wrote the manuscript. AJRB advised and managed HEM in the development of the software and manuscript. DM and SAL contributed to the software under the supervision of KS and BF. DM, SAL, KS, BF, and AJRB critically reviewed and edited the manuscript. All authors read and approved the final manuscript.

## Acknowledgements

We want to thank Lori Kern at Bioconductor for her review of *RLSeq* and *RLHub*. We want to also thank Simon Bray at Bioconda for his review of *RLPipes*. We want to thank the Bioinformatics Research Network for their support in developing *RLSuite*.

## Notes

### Competing Interest Statement

The authors have declared no competing interest.

